# Loss of PMR4 callose synthase results in salicylic acid-independent and broad-spectrum resistance to clubroot in *Arabidopsis* and *Brassica napus*

**DOI:** 10.1101/2024.09.19.613914

**Authors:** Brian Luo, Lipu Wang, Rui Wen, Kun Yang, Xunjia Liu, Jiangying Tu, Tim Dumonceaux, Yangdou Wei, Garry Peng, Wei Xiao

## Abstract

The clubroot disease caused by protist *Plasmodiophora brassicae* is one of the most important diseases of Brassica crops. Growing clubroot-resistant cultivars remains the most effective and practical approach to managing clubroot disease. However, resistance gene-mediated immunity is rapidly overcome in the field when new pathotypes arise. In this study, we identified *PMR4* as a potential gene target for creating a novel clubroot-resistant source. Recessive *PMR4* mutations in *Arabidopsis thaliana* conferred broad-spectrum resistance to multiple *P. brassicae* pathotypes, independent of salicylic acid-mediated plant immunity. CRISPR/Cas9-mediated gene-editing was employed to create mutations in two *PMR4* orthologs in the *B. napus* genome, and resulting homozygous mutants exhibited dual resistance to powdery mildew and clubroot. PMR4 is required for the callose deposition at wound and powdery mildew infection sites in leaves. This study reveals that callose deposition in roots is induced by *P. brassicae* infection and requires PMR4. It appears that the clubroot disease progression is arrested at the primary-to-secondary infection phase in *pmr4-1* mutants. Together, this study demonstrates that *PMR4*-encoded callose synthase is a host susceptibility factor required for *P. brassicae* to complete its life cycle, and that *PMR4* can be targeted against both powdery mildew and clubroot diseases in Brassica crops.

## Introduction

Brassica crops, especially canola/oilseed rape (*B. napus* L.), are cultivated globally with significant economic and nutritional values. However, most of these crops are affected by the clubroot, a severe soil-borne disease caused by the protist *Plasmodiophora brassicae* (*Pb*) (Cardoza and Stewart, 2006; Perez-Lopez et al., 2018). Resting spores, which are the primary inoculum of the pathogen, are capable of surviving for 20 years in the soil (Wallenhammar, 1996). Primary zoospore released from resting spores initially infect root hairs, producing primary plasmodia during this primary infection stage that do not cause macroscopic symptoms (Howard et al., 2010; Hwang et al., 2012). The secondary zoospores released from root hairs penetrate root epidermis of the host and invade cortical tissues of main roots, resulting in club-shaped root symptoms (Hwang et al., 2012). The gall formation impairs water and nutrient uptake, leading to wilting and premature plant death (Hwang et al., 2012; Schwelm et al., 2015).

The most efficient disease management for clubroot to date is growing clubroot-resistant (*CR*) cultivars in various rotations (Hwang et al., 2012; Peng et al., 2015). Although there are several commercial *CR* canola and cabbage cultivars available, this resistance relates to single dominant *CR* genes and can be broken quickly, as observed in Chinese cabbage and oilseed rape, possibly due to diverse and evolving pathogen populations (Matsumoto et al., 2012; Strelkov et al., 2018). Distinct from the *R*-gene strategy, an emerging alternative approach based on susceptibility (*S*) genes has the potential to be more durable in the field than the *R-*gene-driven disease resistance (Zaidi et al., 2018). These *S* genes may make the host compatible with pathogens, facilitating the infection process (van Schie and Takken, 2014). Increasing evidence has shown promise to create novel disease-resistant crops based on the alteration of *S* genes.

With the development of genome-editing techniques like CRISPR/Cas9 in plants (Doudna and Charpentier, 2014; Gao, 2021), the target *S* genes can be readily edited leading to transgene-free products (Soda et al., 2018; Boubakri, 2023) with durable resistance to plant diseases.

We reasoned that *Arabidopsis thaliana* serves as an ideal model system for screening and identification of clubroot *S* (*CS*) genes. Firstly, *A. thaliana* is genetically and physiologically related to the canola/rapeseed species. Secondly, the Arabidopsis ecotype Col-0 is susceptible to all known *Pb* pathotypes. Thirdly, many T-DNA insertion and mutagen-induced mutants in Col-0 are available in the public domain, making the screening of diverse genotypes possible. In this study, we conducted such screening and identified *<u>P</u>OWDERY <u>M</u>ILDEW <u>R</u>ESISTANT 4* (*PMR4*) as a promising *CS* gene. *PMR4* encodes a putative callose synthase (Vogel and Somerville, 2000) and was previously identified as an *S* gene related to the powdery mildew disease on *Arabidopsis*, since loss of the PMR4 callose synthase results in salicylic acid (SA)-dependent plant defense responses against powdery mildew (Vogel and Somerville, 2000; Nishimura et al., 2003). Surprisingly, our experimental data showed that clubroot resistance conferred by the *pmr4* mutation is independent of the SA signaling, favoring a notion that PMR4 serves as a host factor for the pathogenesis of *Pb*. It appears that lack of the PMR4 callose synthase arrests *Pb* zoospore release and progression from primary infection to the secondary infection. Targeted genome editing of *B. napus PMR4* orthologous genes resulted in dual resistance to both powdery mildew and clubroot diseases, making this finding potentially applicable to the Brassica crop improvement.

## Results

### *pmr4-1* confers strong and broad-spectrum resistance against *P. brassicae*

To identify *CS* genes, we developed a strategy to first screen reported plant *S* genes responsible for biotrophic/hemi-biotrophic pathogens and soil-borne diseases, which resulted in over 100 candidate Arabidopsis mutant lines. These mutant plants were inoculated with the *Pb* field isolate 3H, the most prevalent pathotype with high virulence in the western Canada (Hollman et al., 2023), followed by the assessment of clubroot infection and severity at 21 days post-inoculation (dpi). Among 26 mutant lines tested to date (Table S1), a *mlo2,6,12* triple mutant and a *ubc13a,b* double mutant displayed moderate resistance to clubroot. Mildew Resistance Locus O (MLO) proteins are plant-specific calcium channels (Gao et al., 2022) and loss of *MLO* family genes confers resistance to fungal powdery mildew in model plant *Arabidopsis* (Consonni et al., 2006) and crops species such as barley, wheat, pea and tomato (Jorgensen, 1992; Kusch and Panstruga, 2017). *UBC13* encodes ubiquitin conjugation enzymes that mediate non-canonical K63-linked poly-ubiquitination (Hofmann and Pickart, 1999; McKenna et al., 2001). The *Arabidopsis ubc13a,b* double mutation compromises auxin signaling, root growth (Wen et al., 2014) and immune responses (Wang et al., 2019). Surprisingly, a *sweet11,12* double mutant, which was reported to be clubroot resistant (Walerowski et al., 2018) and a corresponding *sweet11,12,15* triple mutant, did not display apparent reduction in the disease severity index (DSI) under our experimental conditions. In sharp contrast, the *pmr4-1* mutant displayed strong resistance with 0% DSI and no obvious galls developed throughout six rounds of experiments (Fig. 1A).

**Figure 1.**
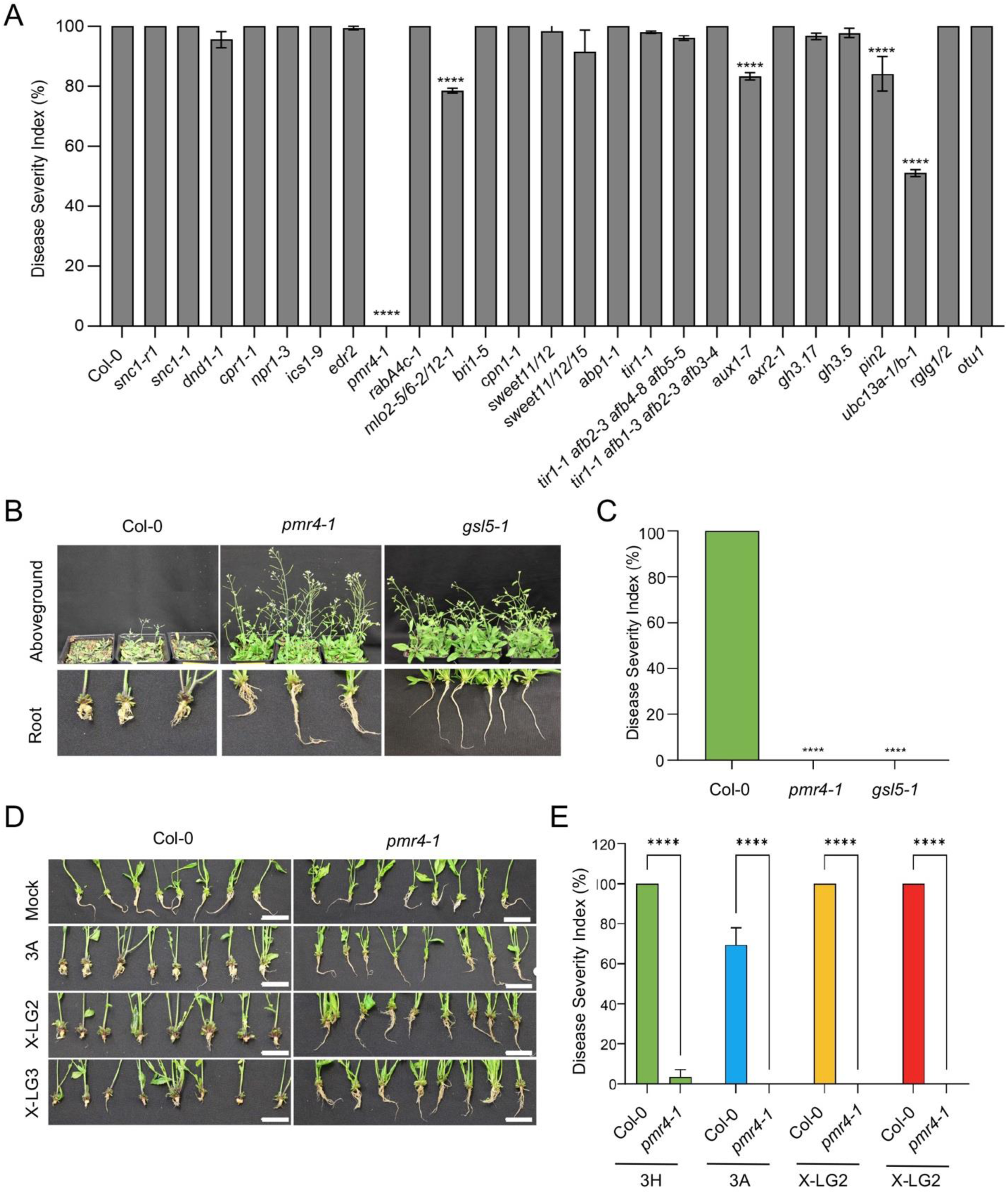
*Arabidopsis pmr4-1* and *gsl5-1* are highly resistance to *P. brassicae* (*Pb*). (**A**) Pathogenicity assays on selected *Arabidopsis* mutants against the clubroot disease. Disease severity index (DSI) reflects quantitative assessment to the disease sensitivity. Clubroot symptoms were rated at 21 dpi after *Pb* pathotype 3H inoculation for at least 4 biological replicates with each lines containing 15 plants. Data are means±SD obtained from 4 independent experiments. ****, P≤0.0001. (**B**) Representative aboveground (upper row) and roots (lower row) images of *Pb* pathotype 3H-inoculated wild type (Col-0), *pmr4-1* and *gsl5-1* mutants captured at 21 dpi. (**C**) Quantitative analysis of *Arabidopsis* clubroot disease severity as shown in (**B**). DSI values are means±SD obtained from four biological repeats with 15 plants for each sample. ****, P≤ 0.0001. (**D**) Representative roots images of Col-0 and *pmr4-1* inoculated with various *Pb* pathotypes captured at 21 dpi. Scale bar = 2 cm. (**E**) DSI (means±SD) of Col-0 and *pmr4-1* inoculated with various *Pb* pathotypes. n=4, ****, P≤ 0.0001.

*PMR4*, also known as *GSL5* and *CalS12*, encodes a predicted trans-membrane callose synthase (Zaveska Drabkova and Honys, 2017). *PMR4* was initially identified because its mutant alleles conferred powdery mildew (PM) resistance (Vogel and Somerville, 2000). The Arabidopsis *pmr4-1* mutation contains a single G-to-A base substitution in the second exon of the *PMR4* gene, resulting in the conversion of a TGG codon for Trp-687 to a TAG stop codon and the formation of a prematurely truncated polypeptide of 686 amino acid residues (Nishimura et al., 2003). This truncated variant of *pmr4* retains the N-terminal six transmembrane helices but lacks the putative catalytic domain and the subsequent seven transmembrane helices at the C-terminus. To independently exam whether the enhanced clubroot resistance is caused by the nonsense mutation in *PMR4,* a T-DNA insertion line *gsl5-1* (GABI-KAT 089H05), in which a T-DNA is inserted to the second exon of *PMR4* close to the nonsense mutation in *pmr4-1* (see Fig. S6) was acquired and inoculated with *Pb* pathotype 3H. Typical swollen root galls were observed in wild-type roots, and the plants were severely affected at 21 dpi, while *pmr4-1* and *gsl5-1* plants remained healthy without visible symptoms both underground and aboveground (Fig. 1B), with DSI values nearly 0% (Fig. 1C). This result indicates that the enhanced clubroot resistance observed in *pmr4-1* and *gsl5-1* mutants was exclusively caused by mutations in the *PMR4* gene.

Since both *pmr4-1* and *gsl5-1* can potentially produce truncated proteins, it is unclear whether they are complete loss-of-function mutations. To determine if the mutations are dominant or recessive for the clubroot resistance, heterozygous *gsl5-1* line and its self-crossing segregants were assessed by a clubroot disease test. The *gsl5-1*/+ heterozygous plants were largely susceptible to the clubroot pathogen, and the F2 progenies segregated 16 resistant and 43 sensitive plants. A chi-square test showed that it fits a phenotypic segregation of 3:1 (χ^2^_3:1_ = 0.142), indicating that the *gsl5-1*-mediated clubroot resistance is recessively inherited. Hence, we conclude that the enhanced clubroot resistance is conferred by a single recessive mutation in the *PMR4/GSL5* gene.

The *pmr4-1* mutant plants were also highly resistant to all tested pathotypes prevalent in Western Canada, including 3H (Fig. 1, B and C), 3A, X-LG2 and X-LG3 (Fig. 1, D and E). In conclusion, loss of *PMR4/GSL5* results in a robust and broad-spectrum clubroot resistance under our experimental conditions.

### *pmr4-1*-conferred clubroot resistance is independent of SA signaling

*pmr4-1* has been well studied in the context of its ability to confer resistance to powdery mildew, a disease caused by fungal pathogens like *Erysiphe cichoracearum* (*Ec*) and *E. orontii*, which infects host leaves and leads to the host cell death (Stone, 1992; Liu et al., 2023). A longstanding explanation for the PM resistance by *pmr4-1* is that callose or callose synthase negatively regulates the SA-mediated immunity. As a result, an enhanced activation of the SA signal transduction pathway is required for the PM resistance in the *pmr4-1* mutant (Nishimura et al., 2003). Given numerous differences between this foliar fungal pathogen and the root-based protist pathogen, we asked whether the enhanced clubroot resistance in the *pmr4-1* mutant is also dependent on the functional SA pathway.

We first tested the SA pathway activation in Col-0 and *pmr4-1* after the *Pb* inoculation, in which relative transcript levels of the *pathogenesis-related 1* (*PR1*) gene was measured by quantitative RT-PCR. Consistent with a previous report (Nishimura et al., 2003), basal levels of *PR1* transcripts in *pmr4-1* were higher in *pmr4-1* than that in Col-0 at all three selected time points (Fig. S1A). Upon *Pb* inoculation, the *PR1* gene were induced at 7, 14 and 21 dpi in Col-0. In comparison, dramatic *PR1* induction was only observed at 7 dpi in *pmr4-1* but not at the other two time points, indicating differential responses of the SA pathway in wild-type and *pmr4-1* following the *Pb* infection.

To determine the SA dependency on *pmr4-1* conferred clubroot resistance, a *pmr4-1 pad4-1* double mutant and a transgenic line *pmr4-1 NahG*, both with compromised SA signaling in the *pmr4-1* mutation background, were subjected to the clubroot disease tests. PAD4 acts upstream of SA accumulation in a positive signal-amplification loop required for the activation of defense responses (Zhou et al., 1998; Jirage et al., 1999), while bacterium-derived NahG specifically degrades SA (Delaney et al., 1994). These lines have been previously used to demonstrate the SA dependency by *pmr4-1* to the PM resistance (Nishimura et al., 2003). To our surprise, a clubroot-resistant phenotype was observed from the *Pb* pathotype 3H-inoculated *pmr4-1 pad4-1* and *pmr4-1 NahG* plants, which were indistinguishable from roots of the corresponding *pmr4-1* single mutant (Fig. 2A). As expected, typical large, swollen root galls were detectable on the Col-0, *NahG*, and *pad4-1* as expected, while healthy roots systems were observed on the *pmr4-1*, *pmr4-1 NahG* and *pmr4-1 pad4-1* plants after the *Pb* inoculation (Fig. 2A). Clubroot DSI further indicates that blocking the SA signaling pathway has no noticeable effect on the *Pb* infection in *pmr4-1* (Fig. 2C). Consistently, *Pb* biomass levels in infected *pmr4-1*, *pmr4-1 NahG* and *pmr4-1 pad4-1* roots remained extremely low and indistinguishable (Fig. 2D), indicating that *pmr4-1* mediated clubroot resistance does not require the SA signaling. Interestingly, *Pb* biomass levels in infected *NahG* and *pad4-1* roots were lower than Col-0 (Fig. 2D), indicating that the SA-mediated immunity plays a minor role in the clubroot resistance.

**Figure 2.**
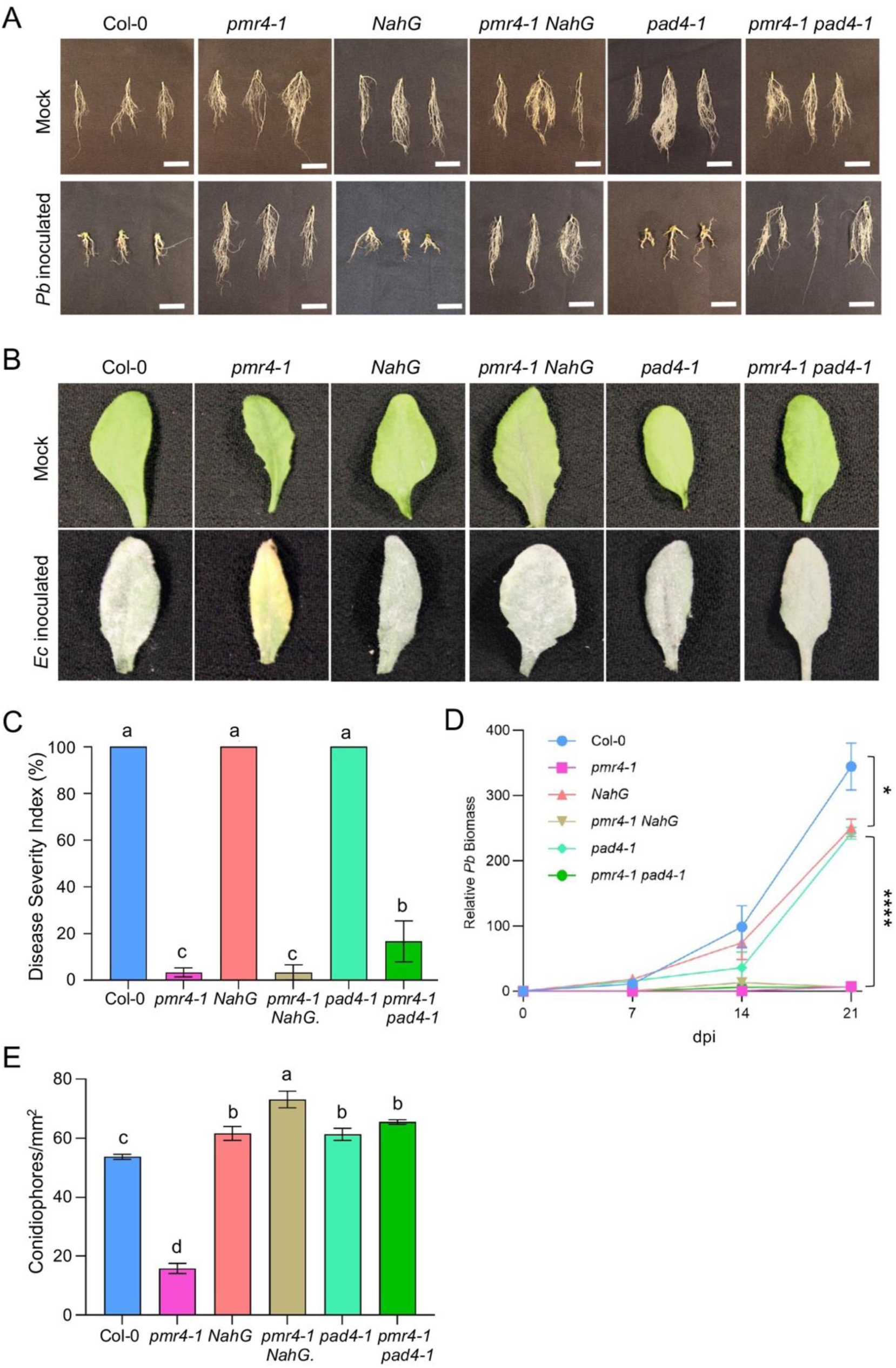
*Arabidopsis pmr4-1* confers dual resistance to *P. brassicae* (*Pb*) and *E. cichoracearum* (*Ec*). (**A**) Representative root images of Col-0 and *pmr4-1* with and without *Pb* pathotype 3H inoculation at 21 dpi. Scale bar = 2 cm. (**B**) Representative leaf images of Col-0 and *pmr4-1* with and without *Ec* inoculation at 15 dpi. Scale bar = 2 cm. (**C**) Quantitative analysis of *Arabidopsis* clubroot disease severity as shown in (**A**). DSI values are means±SD obtained from three independent experiments. One-way ANOVA was performed, and different letters represent various disease severity levels as well as statistical significance. (**D**) Relative *Pb* biomass as measured by qPCR of a *Pb* genomic DNA contents using a genomic DNA segment of the internal transcribed spacer 1 region (Wen et al., 2020). *AtUBQ10* was used as a host tissue normalization reference. At least 3 plant roots were collected as one mixed sample with 3 replicates. Data are means±SD obtained from three technical replicates. One-way ANOVA analysis for *Pb* relative biomass at 21 dpi was performed. *, P≤0.05, ****, P≤0.0001. (**E**) Quantitative analysis of *Ec* conidiophore density on *Ec*-inoculated wild-type and mutant leaves at 5 dpi. One-way ANOVA was performed, and different letters represent statistical significance.

To investigate whether the SA-independency of clubroot resistance in *pmr4-1 pad4-1* and *pmr4-1 NahG* line applies to different *Pb p*athotypes, we inoculated the wild type, *pmr4-1*, *pmr4-1 pad4-1* and *pmr4-1 NahG* plants with *Pb* pathotypes 3A, X-LG2 and X-LG3 *Pb*. Both *pmr4-1 pad4-1* and *pmr4-1 NahG* plants exhibited comparable levels of resistance to various pathotypes as the *pmr4-1* mutant (Fig. S1B). Instead, significant statistical differences were observed between Col-0 and every *pmr4-1* mutant line regardless of their SA status (Fig. S1B).

To further address the distinct requirements of SA defense pathway in *pmr4-1* and determine whether *pmr4-1* can confer dual resistance against PM and clubroot diseases, we simultaneously inoculated the same plants with conidiospores of the host-adapted *Ec* and *Pb* resting spores. As anticipated, typical PM whitish leaves were observed in Col-0, *NahG*, *pad4-1*, *pmr4-1 NahG*, and *pmr4-1 pad4-1*. In contrast, only *pmr4-1* leaves were yellowish, indicative of SA-mediated HR cell death to limit post-penetration invasion (Fig. 2B). In addition, parasitic structures were monitored by acidic aniline blue staining to *Ec*-inoculated leaf samples, which revealed enhanced resistance to *Ec* by the *pmr4-1* mutation, as evidenced by decreased growth, fewer hyphal branches of *Ec* as well as a lower density in conidiophores and disease symptoms on the leaf surface at 7 dpi in *pmr4-1* plants than Col-0 (Fig. S2). Furthermore, inactivation of the SA-signaling pathway in *pmr4-1* mutants led to increased conidiophore growth in *Ec*-inoculated leaves (Fig. 2E), confirming the requirement of SA for *pmr4-1* mediated PM resistance.

Taken together, our results show that although the SA defense signaling is activated in the early stage of *Pb* infection, this signaling pathway plays at most a moderate role in the clubroot resistance phenotype and is not required for the *pmr4-1* conferred broad-spectrum high-level resistance. Importantly, *pmr4-1* exhibited dual resistance to PM and clubroot diseases in SA-dependent and SA-independent manners, respectively.

### Generation of genome-edited *B. napus PMR4* mutants

The *pmr4-1* mutation does not have a significant impact on *Arabidopsis* plant growth and development (Fig. S3), making *PMR4* an ideal target for developing clubroot-resistant crops. Many crops belong to the Brassicaceae family and suffer from the clubroot disease (Hwang et al., 2012). Since loss of the PMR4 activity in Arabidopsis results in clubroot resistance, we asked whether the PMR4 gene is highly conserved in these crops and, if so, whether targeting the PMR4 orthologs in a Brassicaceae crop like *B. napus* also leads to clubroot resistance. The *Arabidopsis* PMR4 amino acid sequence was used to blast the EnsemblPlants database (https://plants.ensembl.org/Multi/Tools/Blast) focusing on Brassicaceae crops with high economic values. A phylogenetic tree based on the candidate PMR4 orthologs retrieved (Fig. S4) indicates that PMR4 orthologs are ubiquitously present and highly conserved in most *Brassicaceae* species. Two genes annotated as *BnaC09g00800D* (*BnaPMR4.C09*) and *BnaA09g01630D* (*BnaPMR4.A09*) were found in the *B. napus* genome, encoding proteins with 91% amino acid sequence identity to *Arabidopsis* PMR4 (E-values at or below 10^−56^) (Fig. S5). Like AtPMR4, predicted BnaPMR4.C09 and BnaPMR4.A09 are polytopic integral membrane proteins with 10 and 12 transmembrane helices, respectively (Fig. S6). In addition, both proteins contain two conserved potential catalytic domains: 1,3-beta-glucan synthase component FKS1-like (FKS1_dom1) and 1,3-beta-glucan synthase component (glucan synthase) (Hsieh et al., 2024).

To create *B. napus PMR4* mutants that mimic Arabidopsis *pmr4-1* and *gsl5-1*, two sgRNA constructs in exon 2 capable of targeting both *PMR4* genes were designed and separately assembled into a CRISPR/Cas9 vector (Fig. 3, A and B). The CRISPR/Cas9 sgRNA constructs were transformed into *B. napus* line DH12075 by using Argobacterium-mediated hypocotyl transformation (Zhou et al., 2002). While no mutation was found in sgRNA4 transformed plants, two sgRNA3 transformed lines were found to carry +1 frameshift mutations in both genes. *pmr4*- *3g37* carries biallelic homozygous (T/T) insertions in *BnaPMR4.C09* (Fig. S7A) and biallelic heterozygous (T/G) insertions in *BnaPMR4.A09* (Fig. S7B). Hence, *pmr4*-*3g37* is a homozygous double mutant for both *BnPMR4* genes in T0. On the other hand, *pmr4*-*3g41* carries monoallelic (T/-) insertions in both *BnPMR4* genes (Fig. S7, A and B), and hence is a heterozygous mutant for both genes in T0.

**Figure 3.**
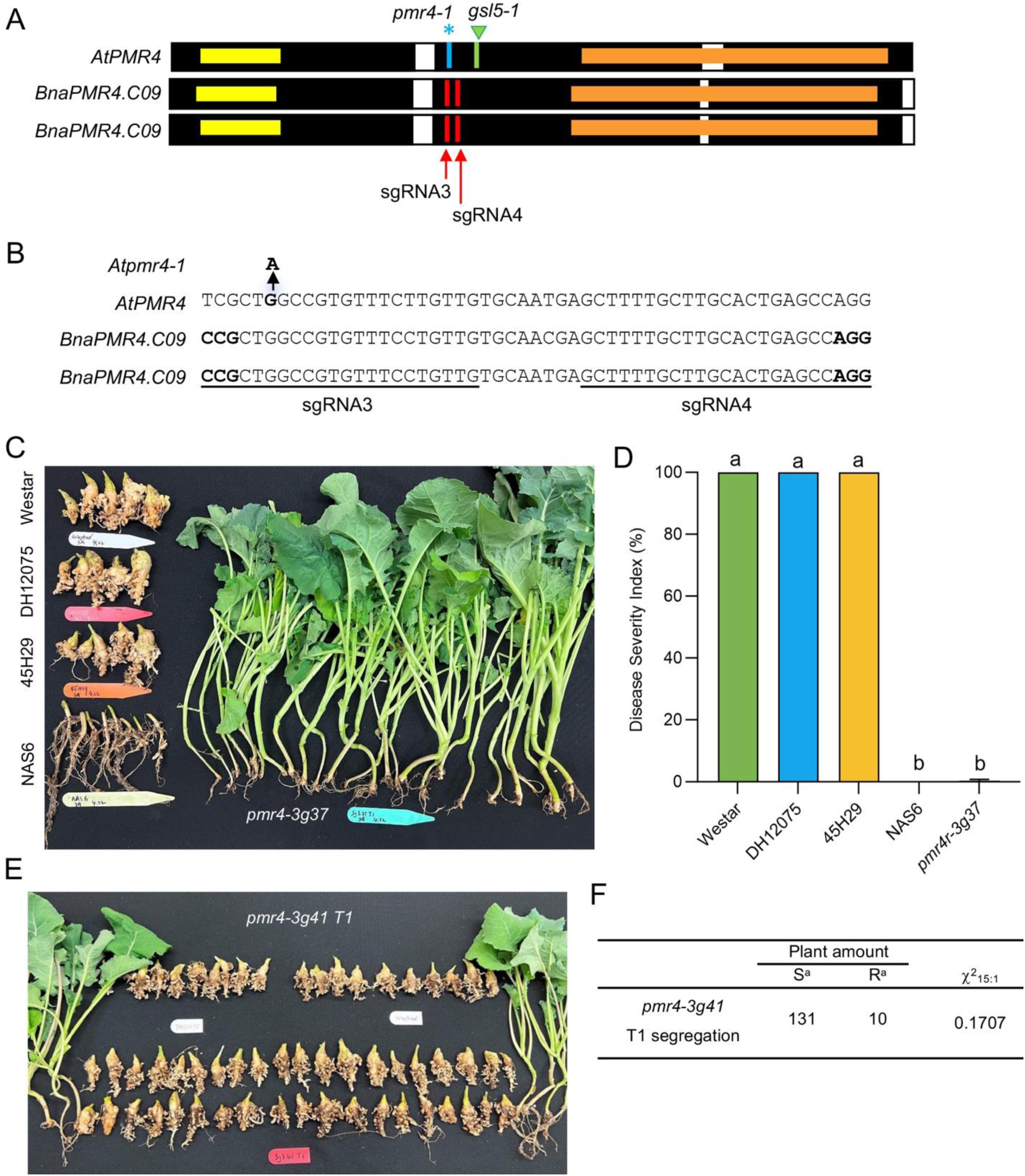
Creation and clubroot disease test of *B. napus pmr4* mutants. (**A**) Comparison of predicted *Arabidopsis* and *B. napus* PMR4 proteins. Filled and open boxes indicate exons and introns, respectively. Yellow boxes represent the FSK1_dom1 domain and orange boxes represent the glucan synthase domain. Blue asterisk indicates the point mutation site in *pmr4-1*; green triangle indicates the T-DNA insertion site in *gsl5-1*; red arrows indicate target sites of single guide (sg) RNAs for *BnaPMR4* genes. (**B**) *B. napus* sequences to show sgRNA target sites. PAM sequences are in bold. (**C**) Representative root and aboveground phenotypes of *B. napus* plants at 28 dpi with *Pb* pathotype 3A. Scale bar = 10 cm. (**D**) Quantitative analysis of results as shown in (**C**). DSI values are means±SD obtained from at least 10 biological repeats. One-way ANOVA was performed, and different letters represent statistical significance. (**E**) Representative root and aboveground phenotypes of *B. napus* heterozygous *pmr4-3g41* self-crossed T1 segregants inoculated with *Pb* pathotype 3A at 28 dpi. (**F**) Summary of phenotypic segregation as shown in (**E**) and χ^2^ test for a two-gene independent segregation hypothesis. S, susceptible plants with disease; R, resistant plants exhibiting no disease symptoms.

### *B. napus PMR4* mutants are clubroot resistant

The homozygous gene edited *pmr4*-*3g37* line was used for the clubroot disease test against *Pb* pathotype 3A along with reference lines including Westar (susceptible commercial line), DH12075 (parental), NAS6 (resistant control) and 45H29 (a previously resistant line that has been overcome by a new *Pb* pathotype) (Jiang et al., 2020). *Pb*-infected *pmr4*-*3g37* plants grew healthy and did not display any chlorosis and stunting phenotypes aboveground (Fig. 3C).

Furthermore, *pmr4*-*3g37* plants formed smaller or no galls on their roots compared to the susceptible lines, which formed large root galls (Fig. 3C). Importantly, *pmr4-3g37* grew normally after the *Pb* inoculation comparable to the *CR* control NAS6, whereas Westar, DH12075 and 45H29 displayed retarded growth (Fig. 3C) and eventually died. Pathotype 3A-inoculated *pmr4-3g37* exhibited a DSI value near 0%, which is comparable to that of the resistant control NAS6 (Fig. 3D). Similar results were also observed in the disease test against pathotype X-LG2 (Fig. S8, A and B), and 3H (Fig. S8, C and D) as *pmr4-3g37* exhibited resistance comparable to corresponding *CR* control.

The heterozygous *pmr4*-*3g41* line was self-crossed and the T1 segregants were subjected to the clubroot disease test against *Pb* pathotype 3A. As shown in Fig. 3E and summarized in Fig. 3F, out of total 141 plants assessed, 10 were clubroot resistant and 131 were susceptible (Fig. 3, E and F), which fits well with the anticipated two-gene segregation ratio of 15:1 if both mutations are recessive. All 10 resistant plants were genotyped and found to contain homozygous mutations in both *PMR4* genes. Similarly, *pmr4*-*3g41* T1 segregants were inoculated with *Pb* pathotype 3H and 8 out 107 plants were resistant (Fig. S8E), all carried homozygous mutations in both *PMR4* genes. The above results collectively demonstrate that *B. napus* homozygous *PMR4* mutants are resistant to various clubroot pathotypes.

### *B. napus PMR4* mutants are resistant to powdery mildew

We also inoculated *pmr4-3g37* plants with host-adapted *Ec* to determine whether *B. napus pmr4* mutants are resistant to PM. Indeed, *pmr4-3g37* plants exhibited strong resistance to PM as canonical whitish powdery phenotypes can be observed on parental DH12075 leaves, while no such disease feature was found from *pmr4-3g37* leaves (Fig. 4A). Surprisingly, the double *R*-gene introgressed clubroot resistant line CPS14 showed susceptibility to the *Ec* infection (Fig. 4A). Meanwhile, microscopic observations of *Ec* parasitic structures after acidic aniline blue staining revealed fewer *Ec* hyphal branches at 7 dpi in *pmr4-3g37* leaves than in Col-0 leaves (Fig. 4B, upper row). Furthermore, *Ec*-inoculated DH12075 and CPS14 leaves at 2 dpi showed characteristic papillae callose deposition, which was dramatically reduced in *pmr4-3g37* leaves (Fig. 4B, lower row), consistent with decreased conidiophore development in *Bnpmr4* mutants (Fig. 4C) and similar to the reported *Atpmr4-1* mutant phenotypes (Jacobs et al., 2003). Hence, we conclude that *BnPMR4* is an ideal target for creating novel *CR* line in *B. napus* that would feature added benefit of PM resistance.

**Figure 4.**
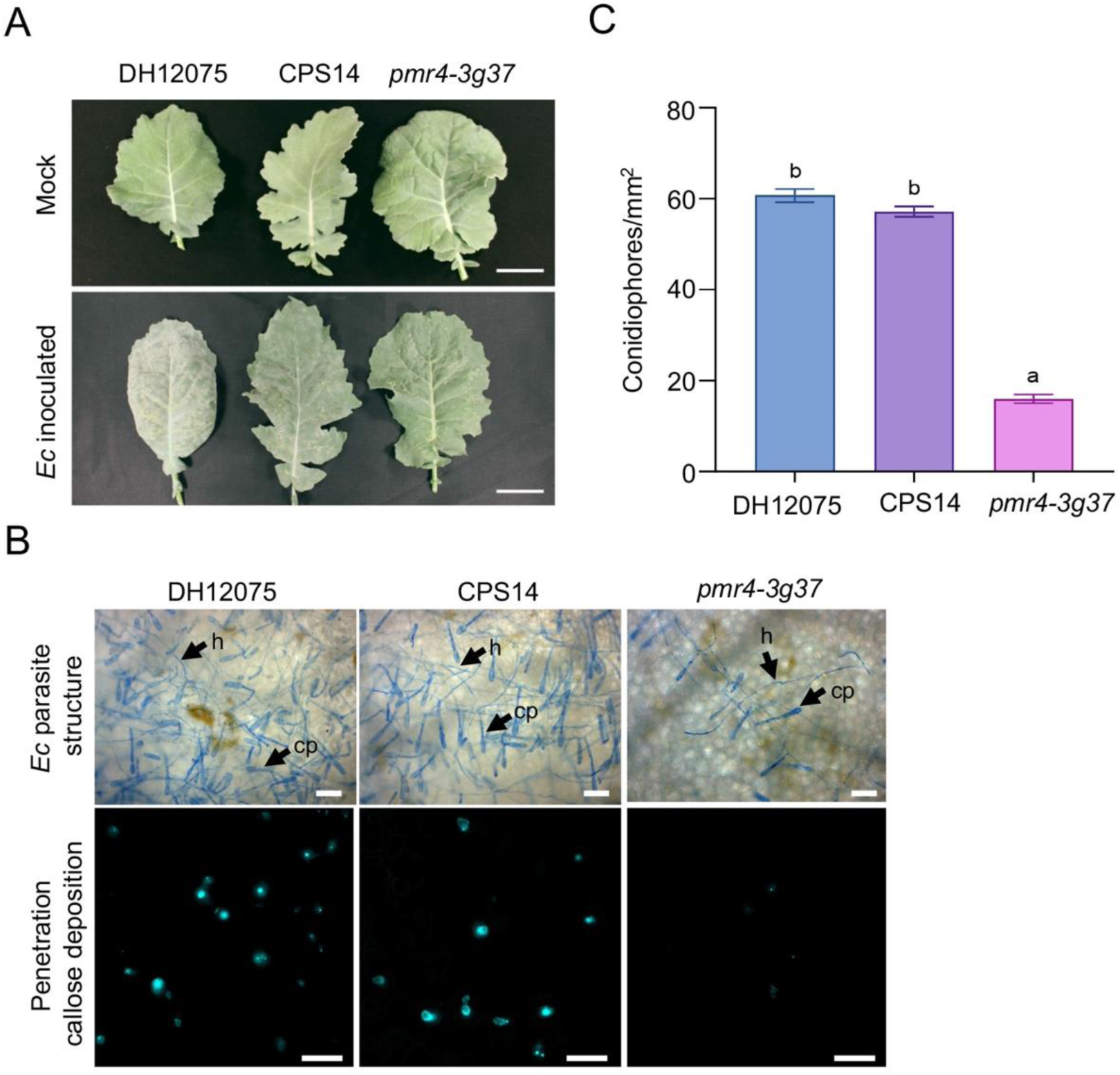
Susceptibility of *B. napus* plants to the powdery mildew disease. (**A**) Leaf phenotypes of DH12075, CPS14 and *pmr4-3g37* 40 days after *Ec* inoculation (lower row). Corresponding uninfected controls are also shown in the upper row. Scale bar = 4 cm. (**B**) Upper row: representative images of hyphae growth and conidiophore production of *Ec* on plant leaf surfaces at 7 dpi. Fungal structures were stained with acidic aniline blue and examined by light macroscopy. h, hyphae; cp, conidiophores. Scale bar = 100 μm. Lower row: callose deposition at the *Ec*-penetration sites stained with basic aniline blue and revealed with epifluorescence microscopy. Scale bar = 50 μm. (**C**) Quantitative measurement of *Ec* conidiophore formation on *Ec*-inoculated leaves at 5 dpi. Data are means±SD from 30 colonies of 3 leaves. One-way ANOVA was performed, and different letters represent statistical significance.

### *P. brassicae* infection leads to PMR4-dependent callose deposition in roots

PMR4 is required for wound-induced callose deposition at the wound sites, which is thought to mimic plant response to the pathogen invasion (Jacobs et al., 2003; Nishimura et al., 2003). A wound-induced callose accumulation test revealed robust callose deposition in Col-0 leaves but is absent in the *Atpmr4-1* leaves (Fig. 5A), which is consistent with previous reports (Jacobs et al., 2003; Nishimura et al., 2003), and hence can serve as positive and negative controls for wounding-induced callose deposition response, respectively. Under the same experimental conditions, DH12075 and heterozygous *pmr4-3g41* T0 leaves displayed clear callose deposition at the wound site 24 hours after the treatment. In contrast, the callose accumulation was absent in both *pmr4-3g37* and homozygous *pmr4-3g41* T1 mutant leaves at the wound site (Fig. 5A). Since the wound-induced callose deposition was still detectable on heterozygous *pmr4-3g41* plant leaves (Fig. 5A), both *pmr4-3* alleles are considered loss-of-function mutations. Hence, PMR4 is required for wound-induced callose deposition in both Arabidopsis and *B. napus*.

**Figure 5.**
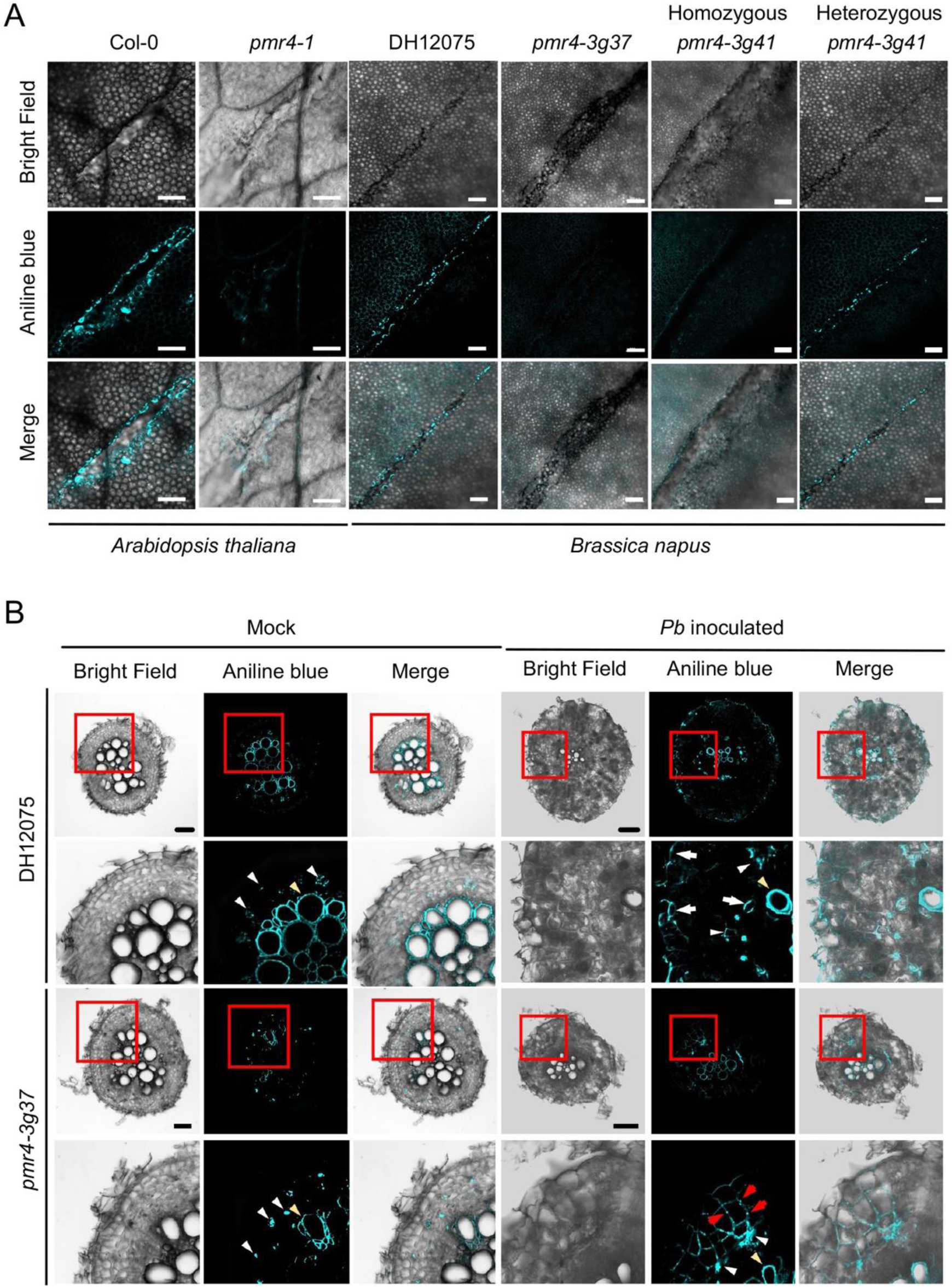
Wound and *P. brassicae*-induced callose deposition in *B. napus*. (**A**) Callose accumulation at wound sites in DH12075 and its *pmr4* mutant derivatives. Arabidopsis Col-0 and its *pmr4-1* mutant serve as references. Plant leaves were wounded by cutting with a razor. The callose accumulation at the wound site was observed 24 hrs after the treatment by aniline blue fluorochrome staining. The heterozygous *pmr4-3g41* plant was from T0, and the homozygous *pmr4-3g41* plant was from genotype-confirmed T1 segregants. Bars = 10 µm. (**B**) Callose deposition patterns of DH12075 and *pmr4-3g37* roots with or without *Pb* pathotype 3H infection. Lateral root samples were collected at 14 dpi, fixed, sliced into 50 μm transverse section and then stained with aniline blue. The boxed region in each upper row was enlarged to form the corresponding lower row. Yellow and white arrowheads indicate xylem lignin auto-fluorescence and sieve element constitutive callose deposition, respectively. White arrows indicate *Pb*-induced callose deposition at epidermal and endodermal cells. Red arrows indicate the punctate PD-localized callose deposition induced. Representative images were selected from samples collected from 5 individual plants, and 3 different sections were observed from each plant. Scale bar = 100 µm.

Taking into account that PMR4 is known as a fungal pathogen and wound responsive callose synthase in leaves (Nishimura et al., 2003), we asked whether the *Pb* infection also induces callose deposition in root tissues and if PMR4 is required for this process. To this end, DH12075 and *pmr4-3g37* plants were inoculated with pathotype 3H resting spores. At 14 dpi, root transverse sections were prepared, and specimens were stained with aniline blue to visualize callose. As shown in Fig. 5B, in the absence of *Pb* inoculation, only sieve element callose (white arrows) and xylem lignin autofluorescence (yellow arrows) were observed in both DH12075 and *pmr4-3g37*, which has been previously reported as a background signal irrelevant to the *Pb* infection (Truernit et al., 2008; Xie et al., 2011). After the *Pb* inoculation, callose accumulation was observed in DH12075 root epidermal and endodermal cells (white arrowheads) but was absent in corresponding *pmr4-3g37* root tissues. Hence, *Pb* infection induces callose deposition in root epidermal and endodermal cells in a PMR4-dependent manner.

Interestingly, a *Pb*-induced punctate callose deposition pattern in cortical cells was observed from *pmr4-3g37* roots (Fig. 5B, red arrowheads) characteristic of a plasmodesmata (PD) callose pattern (Vaten et al., 2011), reminiscent of the callose accumulation pattern on *Ec*-infected *pmr4-1* leaves (Jacobs et al., 2003), suggesting a possibility that PD-callose deposition contributes to the resistance to clubroot and powdery mildew.

### PMR4 is required for the *P. brassicae* secondary infection

To understand why inactivation of PMR4 resists the clubroot disease, we monitored the *Pb* life cycle in Col-0 and *pmr4-1* roots by fluorescent probe-based live cell imaging using a confocal laser-scanning microscopy (CLSM) technique as described (Liu et al., 2020a). Since it has been well established that 7 dpi represents a transition between *Pb* primary and secondary infections, and that the secondary infection phase in cortex starts from 7 dpi and displays disease symptoms at 14-15 dpi (Liu et al., 2020a), we took samples at 7, 14 and 21 dpi to encompass the entire *Pb* life cycle.

The primary infection appeared to be successful and completed in both Col-0 and *pmr4-1* roots since zoosporangia as primary infection parasite structures were found in epidermal cells of both lines at 7 dpi (Fig. 6, A and B). Interestingly, most zoosporangia in *pmr4-1* roots still contained zoospores while only empty zoosporangia could be found in Col-0 roots at 14 or even 21 dpi (Fig. 6, A and B), indicating an inhibition of zoospore release from zoosporangia in *pmr4-1* to prevent further invasion of cortical cells. Consistently, secondary plasmodia and resting spores were found in Col-0 cortical cells at 14 and 21 dpi, indictive of successful secondary infection (Fig. S9). In contrast, few secondary plasmodia were detected in *pmr4-1* root samples, while most zoospores remained unreleased from zoosporangia in root hairs and epidermis (Fig. 6, A and B, Fig. S9), implying that the zoospore release in *pmr4-1* is halted.

**Figure 6.**
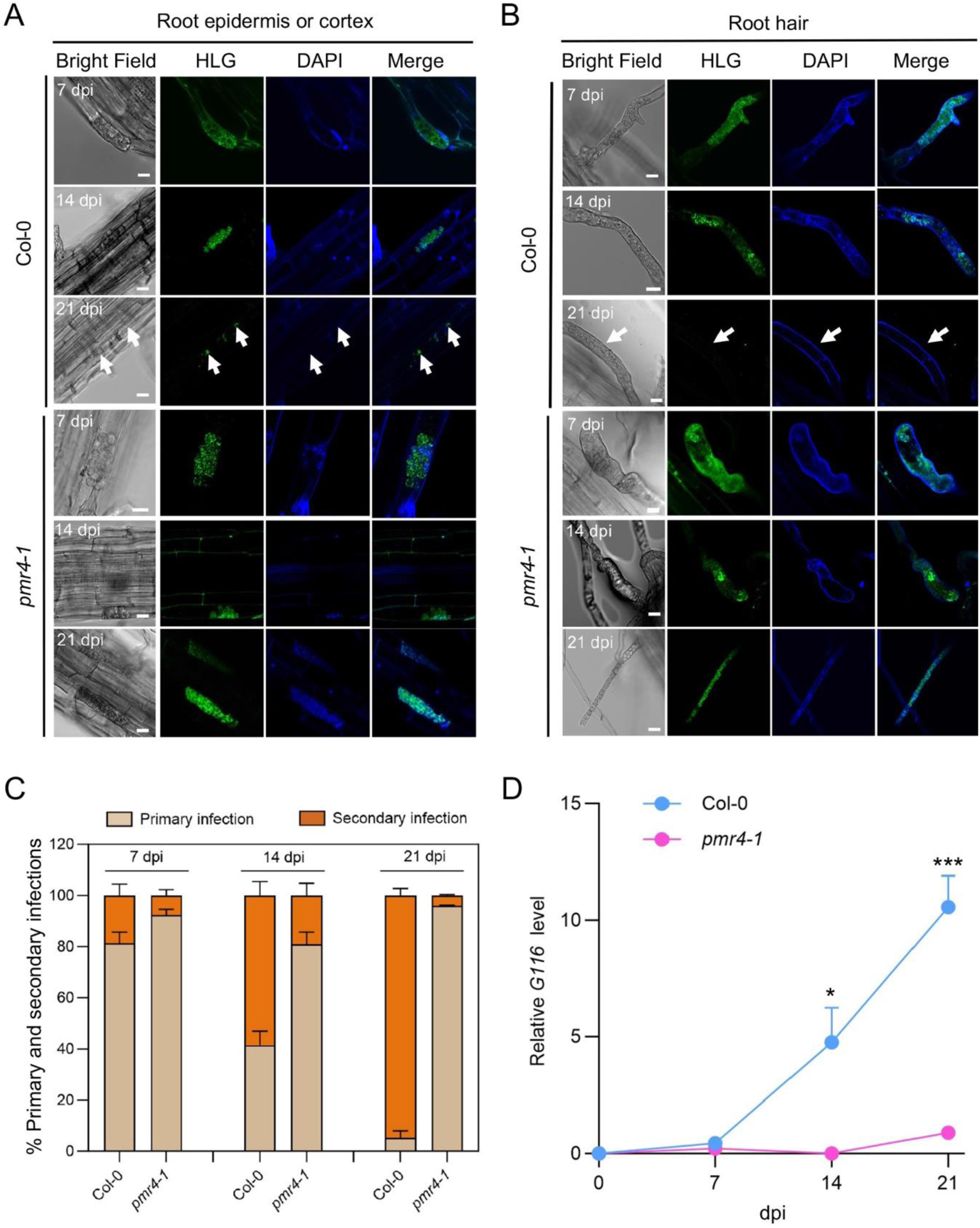
Monitor and quantitative analysis of *P. brassicae* life cycles in *Arabidopsis* roots. (**A**) and (**B**) Confocal laser scanning microscopy images of *Pb* development in root epidermal cells (**A**) and root hairs (**B**). Two-week-old seedlings of Col-0 and *pmr4-1* were inoculated with *Pb* pathotype 3H and roots were collected at 7, 14 and 21 dpi. Live cell dyes HCS LipidTox Green (HLG) and 4’ 6-diamidino-2-phenylinode (DAPI) were used to label *Pb* lipid droplets and nuclei, respectively. Empty zoosporangia were indicated with white arrows. Scale bars = 10 μm. (**C**) Proportions of primary and secondary infection at various time points post inoculation in Col-0 and *pmr4-1*. At each time points except 0 dpi, three 1-cm fragments at the proximal end of the root from three plants were examined to determine the extent of primary and secondary infections using Zeiss LSM880 confocal laser scanning microscopy. On each root fragment, five fields of view were examined using x40 objective lens. Data are means±SD. (**D**) Relative levels of a *Pb* gene *G116* at various time points. *AtUBQ10* was used as a host tissue normalization reference. Data are means±SD and n = 3. *, P ≤ 0.05; ***, P ≤ 0.001, based on two-way ANOVA.

To quantify efficiencies of primary-to-secondary infection transition, we determined proportions of primary and secondary infections at each time point. At 7 dpi, almost all examined root hairs in both lines were infected. At 14 dpi, the primary infection in Col-0 declined to around 40%, while secondary zoospores were released and empty zoosporangia were degraded, accounting for 60% as secondary infections. At 21 dpi, secondary infections in Col-0 increased to over 90%. In contrast, the primary infection accounted for 80-90% at 14 and 21 dpi, while secondary zoospores were rarely observed in *pmr4-1* (Fig. 6C). The relative *Pb* biomass was determined by quantification of *Pb* pathogenesis marker genes like *G116* (Fei et al., 2016). At 7 dpi, the *Pb* biomass was comparable in Col-0 and *pmr4-1*. At 14 and 21 dpi, the relative *Pb* biomass increased dramatically in Col-0 but remained low in *pmr4-1* (Fig. 6D), consistent with a notion that release of most *Pb* zoospores was arrested after the primary infection and hence could not complete their life cycle in *pmr4-1*.

## Discussion

The Brassicaceae family includes many valuable crops, among which canola varieties of *B. napus* are cultivated worldwide for edible oil and other applications due to their high nutritional value and energy outputs. One of the major canola diseases is clubroot caused by a protist *Pb* infection, resulting in devastating yield loss worldwide. Growing *CR* cultivars remains the most effective disease management strategy; however, the molecular nature of existing *CR* genes remains unclear and can be readily broken down due to the pathogen evolution, and *CR* genes were only found in other *Brassica* species that need to be introduced into canola varieties through traditional breeding (Hwang et al., 2012). Recently a novel gene *WeiTsing* (*WTS*) from *Arabidopsis* was identified and characterized. *B. napus* plants carrying *WTS* confer a broad-spectrum clubroot resistance (Wang et al., 2023), making it a promising *CR* gene for the crop improvement. In this study, we developed a strategy to screen *CS* genes from *Arabidopsis*, which can be used to guide the development of clubroot-resistant crops.

Our genetic screen to date has identified several *Arabidopsis* mutant lines displaying moderate clubroot resistance and one mutant line (*pmr4-1*) displaying very strong clubroot resistance. *PMR4* is a previously reported *S* gene for PM (Nishimura et al., 2003). Several pieces of evidence in this study allow us to conclude that *PMR4* is also a *CS* gene. Firstly, two mutant lines carrying independent alleles, *pmr4-1*, containing a premature stop codon in exon 2, and *gsl5-1*, a T-DNA insertion also at exon 2, are resistant to clubroot. Secondly, although it is unclear whether *pmr4-1* and *gsl5-1* are null mutations, they are recessive mutations. Thirdly, the *B. napus* genome contains two copies of *PMR4* orthologs; gene edited mutants become clubroot resistant only when both genes are mutated. Finally, wound and pathogen-induced callose deposition experiments indicate that the above *Arabidopsis* and *B. napus pmr4* mutants are loss-of-function mutations. Although clubroot *S* genes have been previously reported (Chen et al., 2016; Walerowski et al., 2018), to the best of our knowledge, *PMR4* is the first *CS* gene whose inactivation results in such strong clubroot resistance.

Perhaps the most striking finding from this study is that while the PM resistance conferred by *pmr4-1* is dependent on the functional SA pathway (Nishimura et al., 2003), its resistance to clubroot is independent of SA signaling, suggesting mechanistic differences between *pmr4-1* mediated clubroot resistance and that of PM resistance. In the case of PM resistance, it is speculated that either PMR4 synthesized callose deposition in papillae forms a barrier to the fungus as well as the host defense machinery (Jacobs et al., 2003), or this callose negatively regulates the SA signaling (Dong et al., 2008) (Fig. 7A). The SA pathway indeed can play a role in the clubroot resistance under certain circumstances. For example, *bik1*-mediated clubroot resistance is SA-dependent (Chen et al., 2016); a *dnd1* mutation that constitutively activates the immune system alleviates the clubroot symptom (Lovelock et al., 2016); and expression of *WTS* can induce *PR* genes (Wang et al., 2023), although it remains unclear whether *WTS*-mediated clubroot resistance is dependent on the functional SA signaling. Since SA treatment alone can alleviate clubroot symptoms in wild-type plants (Chen et al., 2016), our observed transient induction of *PR* genes in *pmr4-1* upon *Pb* infection may play a role, although it could be undermined by the *pmr4-1* effect.

**Figure 7.**
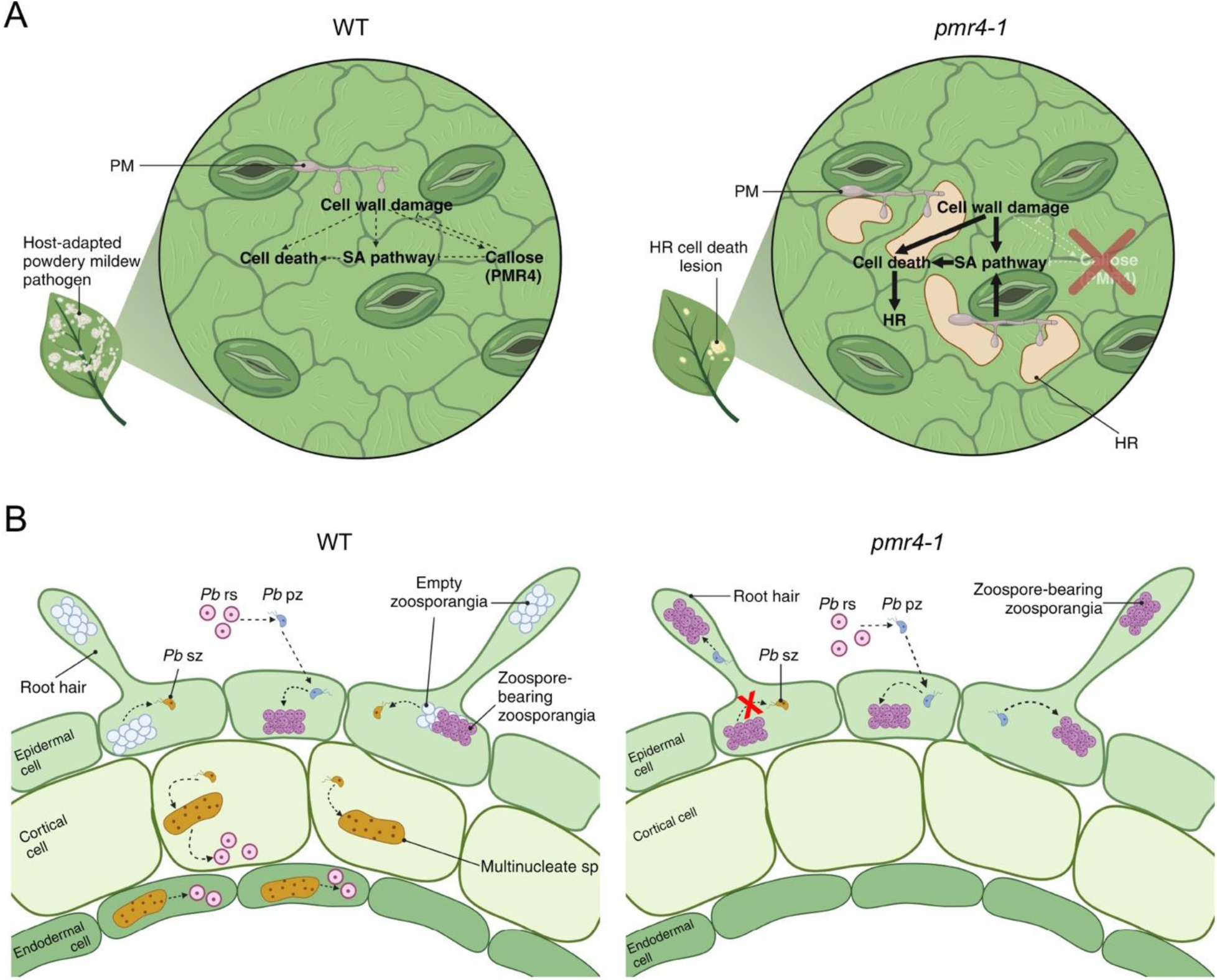
A working model outlining potential mechanisms of *pmr4-1* mediated resistance to powdery mildew (PM) and clubroot in leaves and roots, respectively. (**A**) PM resistance. PM infection damages the cell wall, triggers defense responses such as cell death, SA pathway activation and callose accumulation (cyan color). In *pmr4-1*, because of decrease in the callose deposition at the infection site, the cell wall damage stress cannot be alleviated, and the SA pathway is hyper-activated, resulting in programmed cell death to prevent further development of the pathogen. HR, hypersensitive reaction. Red arrows indicate an induced activation. (**B**) Clubroot resistance. The invasion of *Pb* in wild-type epidermal cells triggers host callose deposition which can be utilized by *Pb* zoosporangia as the source to retrieve the β-1,3-glucan substrate to synthesize their own cell wall. Mature zoospores can be released to keep their virulence. The secondary infection in cortical or endodermal cells results in the abundant production of new resting spores. In *pmr4-1*, although the primary infection successfully occurs, the zoospores cannot be efficiently released due to the lack of host callose supply. rs, resting spores; pz, primary zoospores; sz, secondary zoospores; sp, secondary plasmodia.

Although this study convincingly demonstrated that *pmr4-1* mediated clubroot resistance is independent of SA, it does not rule out the possibility that one of other immunity-related phytohormones, including jasmonic acid (Ghorbel et al., 2021), gibberellins (Verma et al., 2016) or ethylene (Binder, 2020), is required. However, our experimental data favor an alternative hypothesis that PMR4 is a host factor required for the *Pb* life cycle. Firstly, *Pb* can induce callose deposition in *B. napus* root epidermal and endodermal cells, reminiscent of wound and *Ec*-induced callose deposition in *Arabidopsis* leaves (Jacobs et al., 2003). Secondly, the above callose deposition in root tissues is dependent on functional PMR4, indicating that the pathogen utilizes host callose for its infection. Thirdly, the in-house developed CLSM technique and previous observations (Tu et al., 2019; Liu et al., 2020b; Liu et al., 2020a) allowed us to track *Pb* pathogenesis in host roots, which reveals that the release of most zoosporangia in *pmr4-1* was arrested, resulting in delayed or abolished primary-to-secondary infection phase transition and hence the completion of its life cycle. Finally, quantitative analysis of the pathogen load in host roots shows that lack of PMR4 callose synthase prevents the pathogen proliferation and hence the gall formation.

The role of callose deposition in pathogenesis has been the subject of debate (Hsieh et al., 2024). Two hypotheses may be entertained for the PMR4-relatred *Pb* pathogenesis based on literature and our own observations. Firstly, callose is a linear polymer consisting of β-1,3-glucan (Hsieh et al., 2024), which shares the same substrate UDP-glucose with the fungal cell wall polymer β-(1,3; 1,6)-glucan (Onishi et al., 2000; Hu et al., 2023; Hsieh et al., 2024), and hence may be utilized by *Pb* for its cell wall. Secondly, It has been reported that PMR4-independent callose accumulation at PD may restrict the fungal pathogen spread in leaves (Lee et al., 2011; German et al., 2023). For example, a necrotrophic fungus *Rhizoctonia solani* stimulates the expression of *β*-*glucanase* (*OsBGL*) to deposit callose at PD, which reduces its permeability to defend itself against rice sheath blight (Zhou et al., 2023). Similarly, the *Pb*-induced punctate callose pattern in *Bnpmr4* root cortical cells may also alter PD-mediated trafficking, resulting in arrested zoosporangia (Fig. 7B). Future investigation is needed to test these alternative but not necessarily exclusive hypotheses.

Since the *PMR4* gene is highly conserved within the Brassicaceae family, we created the corresponding *pmr4* double mutant lines in *B. napus* by genome editing and demonstrated that they were dually resistant to both clubroot and powdery mildew diseases. Since the PMR4 callose synthase activity is stress induced (Jacobs et al., 2003; Nishimura et al., 2003) and *pmr4-1* mutant plants do not display apparent penalties under normal growth conditions, we conclude that *PMR4* is an ideal target *S* gene for cruciferous crop improvement via gene editing.

## Materials and methods

### Plant materials and growth conditions

Arabidopsis (*Arabidopsis thaliana*) wild type (Columbia-0), and mutant lines used in this study were either purchased from ABRC (https://abrc.osu.edu/) and NASC (https://arabidopsis.info/BasicForm) or kindly provided by other research groups. Detailed information is listed in Supplementary Table 1. All primers used for genotyping and quantitative real-time PCR analysis were listed in Supplementary Table 2. Canola (*B. napus*) cultivar DH12075 was used to generate transgenic *pmr4-3g37* and *pmr4-3g41* plants. For *Arabidopsis* seedling culture, seeds were sterilized and germinated on 1/2 strength Murashige and Skoog (MS) media containing 0.6% agar. For clubroot or powdery mildew disease tests, *Arabidopsis* and *B. napus* seeds were sowed and grown in the soil Sunshine^®^ Mix #5 Natural & Organic (Sun Gro Horticulture Canada Ltd). *Arabidopsis* and *B. napus* plants were grown at 22 °C under 16 h/8 h light/dark photoperiod of ∼125 μE m^-2^ s^-1^ conditions.

### *P. brassicae* and *E. cichoracearum* inoculation

Suspensions of *Pb* pathotypes 3H, 3A, X-LG2 and X-LG3 resting spore inoculum were prepared using galled roots of infected *B. napus* plants. The galled roots were homogenized in a blender with distilled water, and then plant residues were removed from the suspension by filtering through eight layers of cheesecloth. The resting spore concentration was measured by hemocytometer and then diluted to a final concentration at 1 x 10^7^ and 1 x 10^8^ spores per mL for *Arabidospsis* and *B. napus* inoculation, respectively. For *Arabidopsis* inoculation, each 14-day-old seedling was inoculated with 1 mL resting spore inoculum. *B. napus* were inoculated twice with 1 mL spore inoculum each, one at the seeding and another at 7 days after planting when seedlings emerged. For *B. napus* plants the clubroot disease severity was rated using a 0-3 scoring system modified by (Kuginuki et al., 1999). A score of 0 indicates no disease; 1, very small galls on the lateral roots but without deformation on the primary roots; 2, medium galls on the primary roots; 3, severe giant gall formations in the primary roots with or without very little swollen lateral roots left. For *Arabidopsis*, the disease severity was rated using a 0-5 scale as described previously (Hatakeyama et al., 2013) with minor modifications: 0, no visible symptoms; 1, very small galls mainly on lateral roots and that do not impair the main root; 2, small galls covering the main root and few lateral roots; 3, medium to large galls, also including the main root; 4, severe galls on lateral root, main root or rosette; fine roots completely destroyed; and 5, big club on main root and hypocotyl, leading degradation of root; plant growth is affected severely and the plant is dying. The DSI was calculated using the three-grade scale according to the formula:

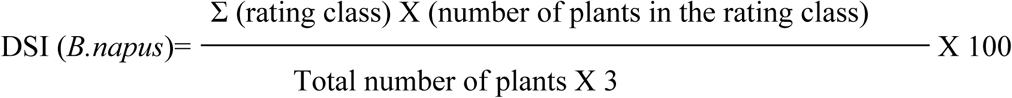

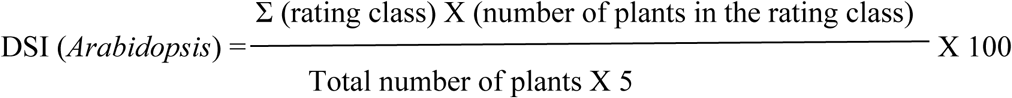

*Ec* was maintained and propagated on *B. napus* plants. 3.5-4.5-week-old *Arabidopsis* plants were inoculated with the conidiospores of *Ec* at a density of 5-10 conidia mm^-2^. For *B. napus*, tested plants were placed in settling towers and inoculated with conidia by heavily tapping the infected *B. napus* leaves above the tested plants.

### Genomic DNA extraction

Root samples were collected and cut by scissors and ground by Lysing Matrix Tubes (FastPrep^®^). The homogenized mixture was used for further genomic DNA extraction by using DNeasy^®^ Plant Mini Kit (Qiagen) following the instruction.

### RNA extraction and RT-qPCR

*Arabidopsis* root samples were collected and rapidly ground in liquid nitrogen into powder, from which 100 mg was used to extract total RNA using the RNeasy^®^ Plant Mini kit (Qiagen). The first-strand cDNA was synthesized from the RNA using iScript™ cDNA Synthesis kit (Bio-Rad). Real-time PCR (qPCR) was conducted against the first-strand cDNA in a real-time PCR detection system using the iQ™ SYBR^®^ Green Supermix and a passive reference dye (Bio-Rad). The qPCR was run in a 20 μL reaction mixture containing 0.5 μL each of 10 μM forward and reverse primers and 50 ng of the cDNA template. The comparative cycle threshold (CT) method was used to analyze and present the data on gene expression according to α ΔΔCT method (Livak and Schmittgen, 2001).

### Confocal laser scanning microscopy

To track *Pb* life cycle root samples were collected at various time points post-inoculation and carefully washed to remove soil residues. Live cell dyes DAPI (Sigma-Aldrich) and HLG (Fisher Scientific) were used to stain the *Pb* parasites following the manufacture’s guide. The DAPI staining was observed by using 405 nm excitation 445 ∼ 480 nm emission wavelengths. The HLG staining was observed by using 488 nm excitation and 499 ∼ 544 nm emission wavelengths. Images were collected with a Carl Zeiss LSM 880 confocal microscope.

To detect *Ec* structures and callose in leaves, treated leaf samples were detached, fixated and destained in 1:3 (v/v) acetic acid/ethanol until the material was transparent (usually overnight). If necessary, the saturated buffer was replaced. Destained leaves were washed in 150 mM K_2_HPO_4_ (pH 9.5) for 30 min or overnight. Leaves were incubated for at least 2 hours in 0.1% aniline blue (2.5% stock solution, Sigma B8563), diluted with 150 mM K_2_HPO_4_ (pH 9.5) in a Falcon tube wrapped in aluminium foil for light protection. 50% glycerol was used for embedding prepared leaf samples in slides. Wound induced callose deposition was observed and collected with Zeiss LSM 880. Confocal images were taken by using 405 nm excitation and 480– 515 nm emission wavelengths to detect callose (cyan).

To detect callose deposition in *Pb-*infected roots, lateral root samples were harvested at 14 dpi and fixed with an FAA solution (37% formaldehyde/glacial acetic acid/95% ethanol/deionized water at a volume ratio of 50:5:10:35) overnight. Samples were embedded in the Tissue Plus O.C.T compound (Leica Microsystems), sectioned with a Leica CM1860 cryostat microtome (Leica Microsystems) to 50 µm and stained with 0.05% ∼ 0.1% aniline blue solution for 15 mins before analysis. After washing three times with ddH_2_O, root sections were subjected to confocal laser scanning microscopy (Zeiss LSM 880). The aniline blue fluorochrome was excited with a 405 nm laser and collected at 480-515 nm.

To observe PM structures, *Ec*-inoculated leaves were detached and collected at different time points, fixed and destained as described above. An acidic aniline blue solution (2 mg/mL, pH 5.0) was used to visualize *Ec* fungal structures using a Zeiss Axioplan 2 upright fluorescence microscope.

### CRISPR/Cas9 vector assembly

Sequence-specific single-guide RNAs (sgRNAs) were designed using the web-based tool CRISPR-P (http://crispr.hzau.edu.cn/cgi-bin/CRISPR2/CRISPR). Target sites were selected based on their location in the target genes, GC% content and putative off-targets. sgRNA-F/R primers (Supplementary Table S2) were annealed and inserted between two *Bsa*I sites of a binary vector pHEE401 obtained from Dr. Qijun Chen (China Agriculture University, China), resulting in pHEE-BnaPMR4-sgRNAs. Plasmid pHEE401 contains a hygromycin resistance marker driven by a cauliflower mosaic virus *35S* promoter, a Cas9 coding sequence driven by an egg cell-specific promoter *EC1.2* and the cloned sgRNA is driven by a *U6-26p* promoter (Wang et al., 2015).

### *B. napus* transformation

The *B. napus* doubled haploid line DH12075 was used as the transformation recipient in this study. The *Agrobacterium tumefaciens*-mediated hypocotyl transformation protocol (Zhou et al., 2002) was followed. Briefly, Agrobacterium strain GV3101 harboring the expression vector was cultured in 20 mL LB liquid medium at 28℃ (180 rpm) for approximately 12 h. When OD600 reached approximately 0.4, 2 mL from culture was aliquoted and centrifuged at 6,000 rpm for 3 mins to collect the pellet. The supernatant was removed, and the pellet rinsed twice with liquid infection medium (LIM; 4.4 g/L MS, 30 g/L sucrose, pH 5.84∼5.88, 200 µM acetosyringone). The resuspended pellet was kept at 4°C for further infection. Sterilized seeds were sowed on 1/2 MS medium and kept in dark for 7 days. Subsequently, hypocotyls were cut to approximately 1 cm into a sterilized plate containing 18 mL pre-added LIM, and then 2 mL of the resuspended pellet was added to the plate and allowed to infect for approximately 10 min. The infected hypocotyls were placed in a cocultivation medium (4.4 g/L MS, 30 g/L sucrose, 18 g/L mannitol, pH 5.84-5.88, 1 mg/L 2,4-D, 0.3 mg/L kinetin, 200 mM acetosyringone and 8 g/L agar) for two days in the dark at 25°C. All hypocotyls were transferred to a calli-inducing medium [CIM, 4.4 g/L MS, 30 g/L sucrose, 18 g/L mannitol, pH 5.84∼5.88, 1 mg/L 2,4-D, 0.3 mg/L kinetin, STS (0.1 M Na_2_S_2_O_3_: 0.1 M AgNO3 = 4:1), 300 mg/L timentin, 25 mg/L hygromycin and 8 g/L agar] for 20 days at 2,000-2,500 lux. The hypocotyls with embryogenic calli were transferred into a shoot-inducing medium (SIM, 4.4 g/LMS, 10 g/L glucose, 0.25 g/L xylose, 0.6 mg/L MES, pH 5.84-5.88, 2.0 mg/L zeatin, 0.1 mg/L IAA, 3 mg/L AgNO_3_, 300 mg/L timentin, 25 mg/L hygromycin and 8 g/L agar) for shoots regeneration, and the medium was replaced every 2–3 weeks until at least 3 leaves appeared. The induced shoots were transferred into a root-inducing medium (RIM, 4.4 g/L MS, 10 g/L sucrose, 300 mg/L timentin and 8 g/L agar, pH 5.84∼5.88). When plantlets grew roots and 4-6 leaves, they were transplanted into pots with nutritive soil and cultivated in a greenhouse.

### Transgenic plant genotyping

Genomic DNA from leaf tissues were used as PCR template to amplify DNA surrounding the CRISPR target sites using specific primers (Supplementary Table 2). The PCR products were directly sequenced, and the sequencing chromatograms were analyzed to identify and distinguish mutated from wild-type sequences adjacent the protospacer adjacent motif (PAM).

## Acknowledgments

We wish to thank Dr. Qijun Chen for the CRISPR cloning vector, Drs. A. Machmair and X, Li for *Arabidopsis* mutants, and Dr. Fengqun Yu (AAFC) for the DH12075 seeds. This work was funded by Natural Science and Engineering Council of Canada Alliance grant ALLRP 576795-22 and Saskatchewan Canola Development Commission/Western Grains Research Foundation (WGRF) grant ADF 20210893 to W.X.

## Author contributions

L.W. developed the mutant screen strategy; B.L. and R.W. conducted the mutant screen. B.L. characterized *Arabidopsis pmr4* mutant phenotypes under supervisions of J.T. and Y.W. R.W., K.Y. and X.L. created and characterized *B. napus pmr4* mutants under supervisions of T.D. and G.P. L.W. and W.X. designed and supervised overall experiments; B.L., L.W. and W.X. wrote the manuscript with inputs from all co-authors. The authors declare no conflict of interest.

## Supplementary materials

Table S1. *Arabidopsis* mutant lines used in this study. Table S2. Oligonucleotides used in this study.

Fig. S1. Genetic relationship between *Arabidopsis PMR4* and the SA pathway

Fig. S2. Microscopic analysis of the *Ec* pathogen in infected *Arabidopsis* leaves

Fig. S3. *pmr4-1* life cycle in comparison to wild-type Col-0 under normal growth conditions

Fig. S4. A phylogenetic tree of *PMR4/CalS12* orthologs in representative Brassicaceae crops

Fig. S5. Amino acid sequence alignment of AtPMR4 and its two orthologs identified from the *B. napus* genome

Fig. S6. Predicted AtPMR4, BnaPMR4.C09 and BnaPMR4.A09 protein topology

Fig. S7. Histograms showing nucleotide sequences of *pmr43g37* and *pmr4-3g41* T0 at *BnaPMR4* loci surrounding the sgRNA3 PAM sequence

Fig. S8. Assessment of *B. napus pmr4* mutants to the clubroot susceptibility infected by different *Pb* pathotypes

Fig. S9. Assessment of *Pb* secondary infection in Col-0 and *pmr4-1*.

